# Protein language model embeddings for fast, accurate, alignment-free protein structure prediction

**DOI:** 10.1101/2021.07.31.454572

**Authors:** Konstantin Weißenow, Michael Heinzinger, Burkhard Rost

## Abstract

All state-of-the-art (SOTA) protein structure predictions rely on evolutionary information captured in multiple sequence alignments (MSAs), primarily on evolutionary couplings (co-evolution). Such information is not available for all proteins and is computationally expensive to generate. Prediction models based on Artificial Intelligence (AI) using only single sequences as input are easier and cheaper but perform so poorly that speed becomes irrelevant. Here, we described the first competitive AI solution exclusively inputting embeddings extracted from pre-trained protein Language Models (pLMs), namely from the transformer pLM ProtT5, from single sequences into a relatively shallow (few free parameters) convolutional neural network (CNN) trained on inter-residue distances, i.e. protein structure in 2D. The major advance originated from processing the attention heads learned by ProtT5. Although these models required at no point any MSA, they matched the performance of methods relying on co-evolution. Although not reaching the very top, our lean approach came close at substantially lower costs thereby speeding up development and each future prediction. By generating protein-specific rather than family-averaged predictions, these new solutions could distinguish between structural features differentiating members of the same family of proteins with similar structure predicted alike by all other top methods.

## Introduction

### Protein structure prediction problem solved

The bi-annual meeting for Critical Assessment of protein Structure Prediction (CASP) has been serving as a gold-standard for the evaluation of protein structure prediction for almost three decades (Moult et al., 1995). At its first meeting (CASP1 Dec. 1994), the combination of machine learning (ML) and evolutionary information derived from multiple sequence alignments (MSAs) reported a major breakthrough in secondary structure prediction (Rost & Sander, 1995). This concept, expanded into deep learning inter-residue distances (Jones et al., 2015; Li et al., 2021; Wang et al., 2017) which through Alphafold’s deep dilated residual network became so accurate to serve as constraints for subsequent folding pipelines (Kryshtafovych et al., 2019; Senior et al., 2020). Now, DeepMind’s *<u>AlphaFold 2</u>* (Jumper et al., 2021) has combined more advanced artificial intelligence (AI) with larger and more complex MSAs to essentially solve the protein structure prediction problem: at least in principle, predictions now can directly support experimental structure determination (Flower & Hurley, 2021). However, even this pinnacle of 50 years of research has two major shortcomings: (i) predictions are more family-specific than protein-specific, (ii) structure prediction requires substantial computing resources, although 3D structure predictions have been made available for 20 entire proteomes with more to come soon (Tunyasuvunakool et al., 2021).

All competitive structure prediction methods, including *AlphaFold 2*, rely on correlated mutations (Marks et al., 2011). Direct Coupling Analysis (DCA) sharpens this signal (Anishchenko et al., 2017) either through pseudolikelihood maximization (Balakrishnan et al., 2011; Seemayer et al., 2014) or through sparse inverse covariance estimation (Jones et al., 2011). This fails for MSAs lacking diversity (too little signal) and is challenging for families with too much diversity (too much noise). The differentiation of co-evolving residue pairs arising from intra-protein contacts or inter-protein interactions further complicates *de novo* structure prediction (Uguzzoni et al., 2017). One solution is to generate multiple MSAs with different parameters, alignment tools and databases (Jain et al., 2021; Zhang et al., 2020), rendering the input generation even more time-consuming, e.g. 212 minutes for our test sets of 31 proteins on an Intel Xeon Gold 6248 (Results).

### Protein language models (pLMs) decode aspects of the language of life

In analogy to the recent leaps in Natural Language Processing (NLP), protein language models (pLMs) learn to “predict” masked amino acids given their context using no other annotation than the amino acid sequences of 10^7-10^9 proteins (Alley et al., 2019; Asgari & Mofrad, 2015; Bepler & Berger, 2019, 2021; Elnaggar et al., 2021; Heinzinger et al., 2019; Madani et al., 2020; Ofer et al., 2021; Rao et al., 2019; Rives et al., 2021; Wu et al., 2021). Toward this end, NLP words/tokens correspond to amino acids, while sentences correspond to full-length proteins in the current pLMs. <u>Embeddings</u> extract the information learned by the pLMs. In analogy to LMs in NLP implicitly learning grammar, pLM embeddings decode some aspects of the language of life as written in protein sequences (Heinzinger et al., 2019; Ofer et al., 2021) which suffices as exclusive input to many methods predicting aspects of protein structure and function without any further optimization of the pLM using a second step of supervised training (Alley et al., 2019; Asgari & Mofrad, 2015; Elnaggar et al., 2021; Heinzinger et al., 2019; Madani et al., 2020; Rao et al., 2019; Rives et al., 2021), or by refining the pLM through another supervised task (Bepler & Berger, 2019, 2021; Littmann, Heinzinger, et al., 2021). Embeddings can outperform homology-based inference based on the traditional sequence comparisons optimized over five decades (Littmann, Bordin, et al., 2021; Littmann, Heinzinger, et al., 2021). With little additional optimization, methods using only embeddings without any MSA even outperform advanced methods relying on MSAs (Elnaggar et al., 2021; Stärk et al., 2021). In the simplest form, *<u>embeddings</u>* mirror the last “hidden” states/values of pLMs. Slightly more advanced are weights learned by a particular type of LM, namely by transformers; in NLP jargon, these weights are referred to as the “*<u>attention heads</u>*” (Vaswani et al., 2017). They directly capture complex information about protein structure without any supervision (Rao et al., 2020) relating to Potts-models (Bhattacharya et al., 2020). Transformer models can also process MSAs to improve predictions (Jumper et al., 2020; Rao et al., 2021), an advantage at the price of the aforementioned issues with MSA-based predictions.

Here, we introduced a novel approach toward using attention heads (Ahs) from pre-trained transformer pLMs to predict inter-residue distances without MSAs at levels of performance similar to top methods relying on large MSAs and evolutionary couplings/DCA. Thereby, this approach enables accurate predictions of protein 3D structure substantially faster and at lower computing costs.

## Methods

### Data set

We obtained 77,864 high-resolution experimental three-dimensional (3D) structures from ProteinNet12 (AlQuraishi, 2019) compiled from the PDB (Burley et al., 2017) before the CASP12 submission deadline (Moult et al., 2018) thereby replicating the CASP12 conditions. To save energy, we trained on a redundancy-reduced dataset by selecting cluster representatives using MMseqs2 (Steinegger & Söding, 2017) at 20% pairwise sequence identity (PIDE), ultimately training on 21,240 of the 77,864 proteins (SetTrnProtNet12).

ProteinNet12 included a validation set with 41 protein chains from the CASP12 targets for model optimization (SetValCASP12). We used the free-modeling and so called “template-based modeling-hard” (TBM-hard) targets from CASP13 (Kryshtafovych et al., 2019) and CASP14 with publicly available experimental structures (15 for CASP13: SetTstCASP13, 16 for CASP14: SetTstCASP14) as test sets to assess performance.

For the baseline model comparison, we used an in-house model trained on co-evolution/evolutionary couplings. We used the MSAs provided by ProteinNet12 and generated alignments for our additional CASP test sets using the EVcouplings webserver (Hopf et al., 2019) on UniRef100 (Suzek et al., 2015) with bitscore thresholds between 0.1 and 0.7. CCMpred (Seemayer et al., 2014) optimized Potts model hyperparameters.

### Input

As input for the prediction of inter-residue distances, we compared two different types of hidden states derived from pre-trained pLMs: (1) The hidden state output by the last layer of the pLM (for SeqVec (Heinzinger et al., 2019) the last LSTM layer; for the transformer-based models, ProtBert, ProtAlbert, and ProtT5 (Elnaggar et al., 2021), the last attention layer), or (2) the attention scores of each of the attention heads (Ahs) of transformers (not for SeqVec). The advantage of the latter is that we make use of the attention’s all-against-all comparisons between all tokens/residues in sentence/sequence which automatically results in a L-by-L representation for a sequence of length L. As detailed elsewhere (Elnaggar et al., 2021), we used only the Encoder-part of the ProtT5 and created embeddings in half-precision mode to speed-up the embedding generation.

When training on attention heads extracted from ProtT5, the resulting pairwise tensors of shape LxLx768 (24 attention layers, each with 32 attention heads resulting in a total of 768 attention score matrices) would require immense memory and substantially increase training time. To save resources, we trained a logistic regression (LR) model on 200 randomly selected samples from our training set to predict distance probability distributions, evaluated performance on medium- and long-range contact performance for the CASP12 validation set and selected the Top-50, Top-100 and Top-120 attention heads based on the absolute value of the learned weights of the LR. As attention scores may be asymmetric, we enforced symmetry by applying average product correction (APC) as suggested previously (Rao et al., 2020). For each attention head of shape LxL, we computed the APC as follows:

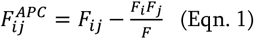

where *F*_*i*_ is the sum over the i-th row, *F*_*j*_ is the sum over the j-th column and *F* the sum over the full matrix.

### Model architecture

Irrespective of the input, our deep learning (DL) models consisted of deep dilated residual networks similar to *AlphaFold 1* (Senior et al., 2020). Each residual block consisted of three consecutive layers (Fig. 1): (1) a convolution with kernel size 1 reduced the number of feature channels from 128 used in the residual connections to 64, (2) a dilated convolution with kernel size 3 (Yu & Koltun, 2015), and (3) a convolution scaling the number of feature channels back up to 128. The dilation factor cycled between 1, 2, 4 and 8 in successive residual blocks. In each layer, we used batch normalization, followed by exponential linear units (ELU) for non-linearity (Fig. 1). Expecting the optimal number of residual blocks necessary to vary for different inputs, we tried depths between 4 and 220 blocks.

**Fig. 1:**
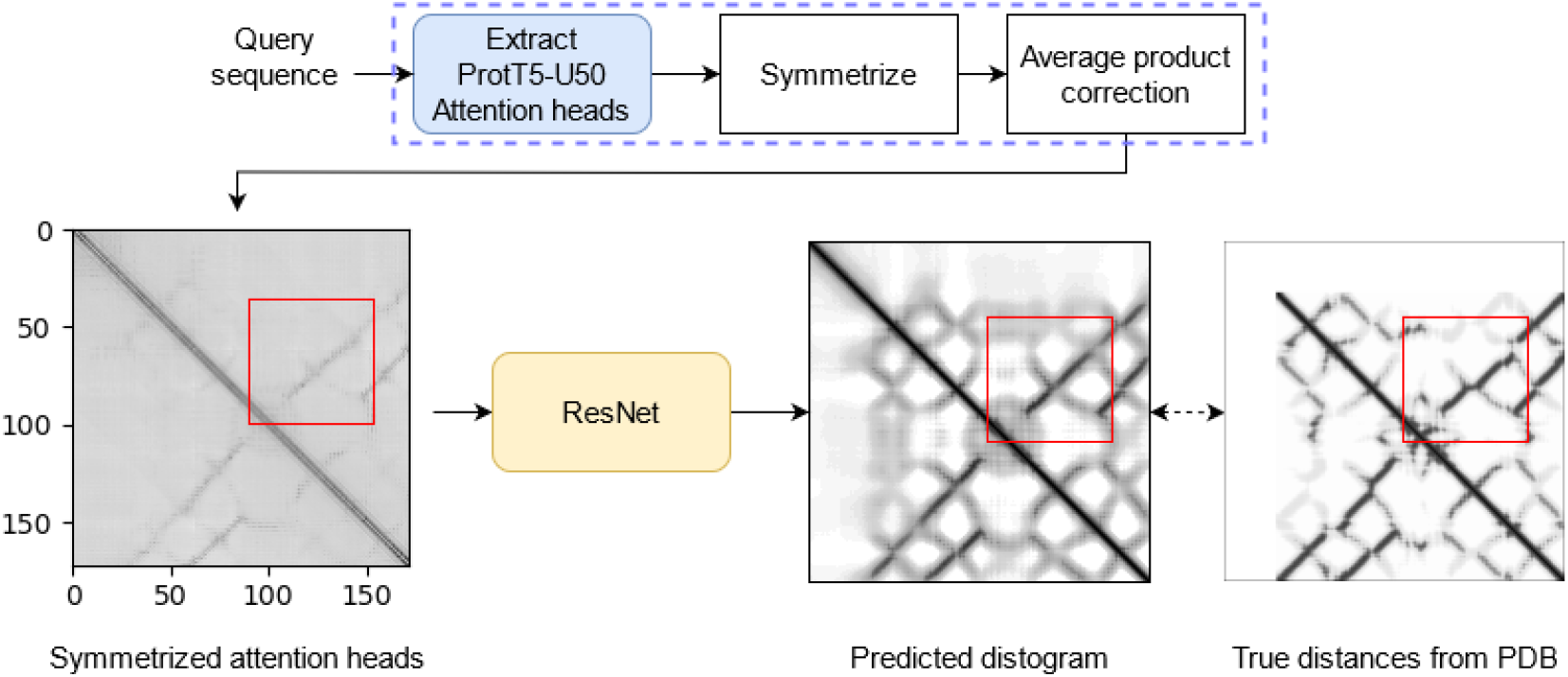
Sketch of approach. The residual CNN (ResNet, yellow-orange) is similar across models (Fig. SOM_1). Models using 1D protein embeddings from ProtT5 (shown here), ProtBERT, ProtAlbert, or SeqVec adapted their architecture to account for expansion from 1D to 2D (Methods). The red square illustrates that proteins were split into overlapping crops of size 64×64 (64 consecutive residues) as introduced by *AlphaFold 1* (Senior et al., 2020). For each crop the CNN (ResNet) predicted the corresponding patch of the distogram. The example above shows the average over the attention heads after applying symmetry and APC for T1026 (CASP14), suggesting that ProtT5 already learned aspects of protein structure without supervision. Our baseline model trained on information from co-evolution for comparison replaced the modules marked by a dashed blue line by the generation of MSAs and estimation of parameters for the Potts model.

Inputting co-evolution information, a narrow window/square around a pair of residues suffices to correctly infer contacts (Jones & Kandathil, 2018). As *AlphaFold 1*, we addressed this through <u>cropping</u>, i.e. by training and evaluating on patches of 64×64 residue pairs extracted from the full distance map.

The two different types of input, 1D protein embeddings (string of numbers) and 2D attention heads (matrix of probabilities), required two different architectures. The architecture predicting distances from 2D attention heads resembled *AlphaFold 1* (Fig. SOM_1), that inputting 1D protein embeddings accounted for the change in input shapes as follows (A) The architecture for 1D embeddings used residual blocks (Fig. 1, Fig. SOM_1), with 1D convolutions for the first half of all residual blocks (e.g. for 120 blocks, 60 residual blocks were 1D convolutions, another 60 blocks were 2D convolutions). Between the 1D and 2D parts, the 1D representations with length L were expanded to pairwise representations of shape LxL. (B) Our models inferred a distance probability distribution (distogram) over 42 bins representing distance intervals between 0-22 Ångstrøm (2.2 nm). The 40 central bins represented distance intervals of 0.5 Ångstrøm, the first 0.2nm (0-2 Ångstrøm) and the last everything else (>22 Ångstrøm). To also assess the performance in predicting contacts rather than distances, we summed the predicted probabilities of the first 14 bins representing distances below 8 Ångstrøm (Wang et al., 2017).

We trained deep learning systems on protein embeddings from a variety of pLMs as well as on co-evolution inputs for baseline comparison. We confirmed 220 residual blocks as optimal when using co-evolution as input (Senior et al., 2020). Seqvec and ProtAlbert embeddings already reached their peak performance with 110 blocks, while ProtBert-BFD (subsequently referred to as ProtBert) required 220. Using ProtT5-U50 (subsequently referred to as ProtT5) we could already reach peak performance with 80 blocks (Table 2), both for embeddings and attention head inputs respectively.

We trained on non-overlapping crops, including patches up to 32 residues off-edge with zero-padding and masking at the edges. To avoid introducing bias by similar structural motifs in the protein ends, we randomly picked the initial offsets for each training sample between -32 and 0 (Senior et al., 2020).

We used overlapping crops with a stride of 32 for training and evaluation (cross-training, i.e. hyper-parameter optimization) and 16 for the final inference for testing/performance estimation. As the number of strides inversely correlated with compute time, this sped up training while providing more reliable predictions at the end. Predictions for residue pairs were averaged across patches to obtain full distance maps. Since distances near the center of each crop were predicted better (more local information available), we computed the weighted average of overlapping predictions by using a Gaussian kernel, giving higher emphasis to central pairs.

### Training

We trained using the Adamax optimizer with an initial learning rate of 1e-2 and a batch size of 75. We performed early stopping and saved the best model checkpoint when the MCC on our validation set (CASP12) did not improve over ten iterations.

### Input

The main input features for our models were either protein representations derived from our pLMs or, for comparison to a baseline, the co-evolution signal in the form of Potts model parameters. To both, we added normalized residue positions (relative position in the protein between 0 and 1), normalized protein length and the log-normalized number of effective sequences as additional input channels. We also masked residues not resolved experimentally, both as single amino acid input and as residue pair during the loss computation.

### 3D predictions

We used pyRosetta (Chaudhury et al., 2010) to compute 3D structures by using a modified version of the trRosetta folding protocol (Yang et al., 2020). In contrast to trRosetta, we dropped any constraints on angular information and we adapted the script such that C-alpha instead of C-beta distances were used as constraints. We first generated 150 coarse-grained decoys using short-, mid- and long-range distances from our predicted distograms at varying levels of distance probability thresholds (here: [0.05, 0.5]) as constraints and relaxed 50 models by using pyRosetta’s FastRelax protocol. Decoy selection as well as the selection of the final model were based on the lowest total energy reported by Rosetta.

For comparison with state-of-the-art (SOTA) methods using MSAs, we obtained 3D models and distance predictions for our test sets (SetTstCASP13 + SetTstCASP14) from the Raptor-X web server (Wang et al., 2017) (accessed June 2021). We only submitted the original query sequences instead of MSAs to allow Raptor-X to follow its own protocol.

### Performance measures

We used performance metrics established by CASP to evaluate the performance of our models, including precision (Eqn. 2), recall (Eqn. 3), F1-score (Eqn. 4), Matthew’s correlation coefficient (MCC, Eqn. 5) and Top-L precision, which measures the positive predictive value of the L long-range contacts predicted with the highest probability (L representing protein length). Specifically, we reported the performance on the L/1, L/2, L/5 and L/10 residue pairs per protein. We adopted the common thresholds of >4 and >23 residues sequence separation to define medium- and long-range contacts respectively and omitted evaluating short-range contacts (|i-j|≤4).

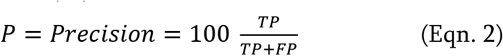

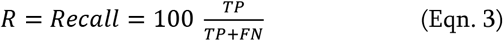

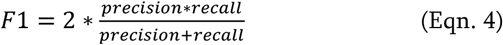

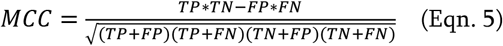

3D predictions resulting from predicted distances, were assessed through TM-align (Zhang & Skolnick, 2005).

### Error estimates

We computed the standard error as usual:

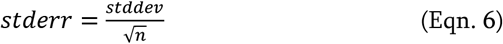

With n as the number of proteins, and stddev the standard deviation obtained by NumPy (Harris et al., 2020). We reported the 95% confidence interval (CI95), i.e. 1.96 standard errors in results:

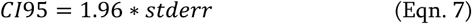

## Results & Discussion

### Top 100 attention heads (AHs) almost as good as all 768 at lower costs

A <u>Logistic Regression</u> (<u>LR</u>) using a subset of the training set and only the validation set (*SetValCASP12*) for optimization suggested that about a one-seventh of all 768 <u>attention heads (AHs</u>) sufficed to get close to saturation in performance although using 7-times fewer parameters (Table 1). This reduced the total storage requirement for training (3.1 TB to 406 GB), in turn enabling local storage for faster data loading, thereby speeding up. That a model as simple as the LR sufficed, highlighted the remarkably strong structural signal readily available from the attention heads of ProtT5 (Elnaggar et al., 2021). Although trained only on a minute set of 200 proteins (100-fold smaller than the 21,240 in the training *SetTrnProtNet12*), the resulting model outperformed convolutional neural networks (CNNs) completely trained on less complex embeddings (Seqvec (Heinzinger et al., 2019) and ProtAlbert (Elnaggar et al., 2021); Fig. 2B).

**Table 1:**
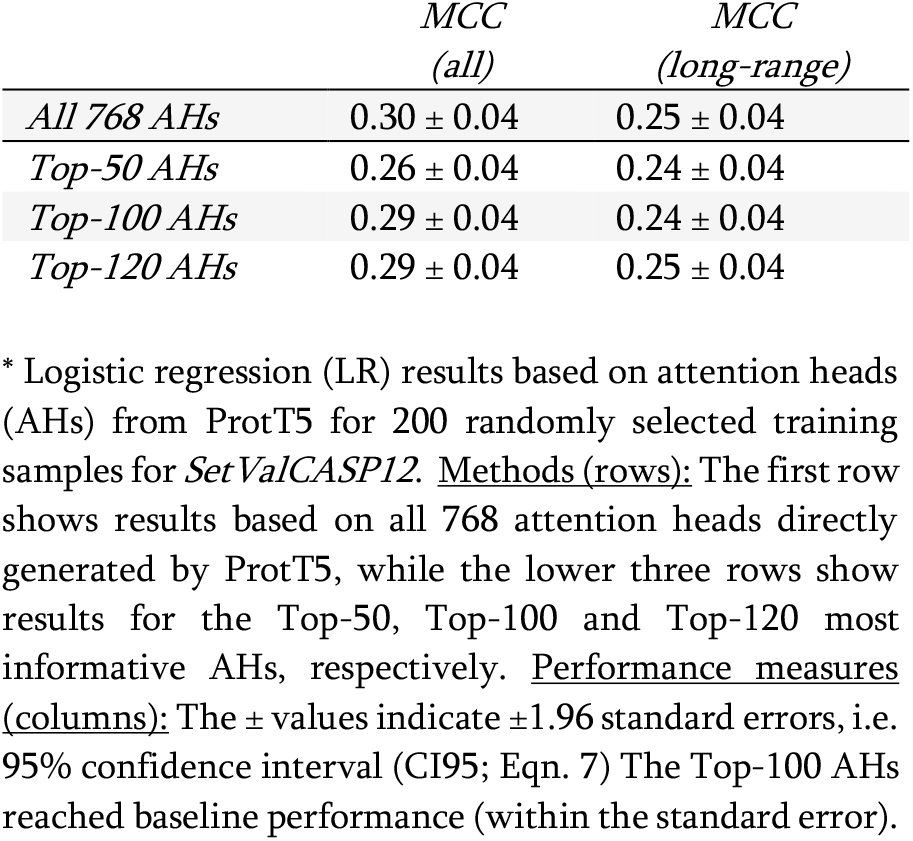
**Performance saturation reached for subset of attention heads (AHs)\***

**Fig. 2:**
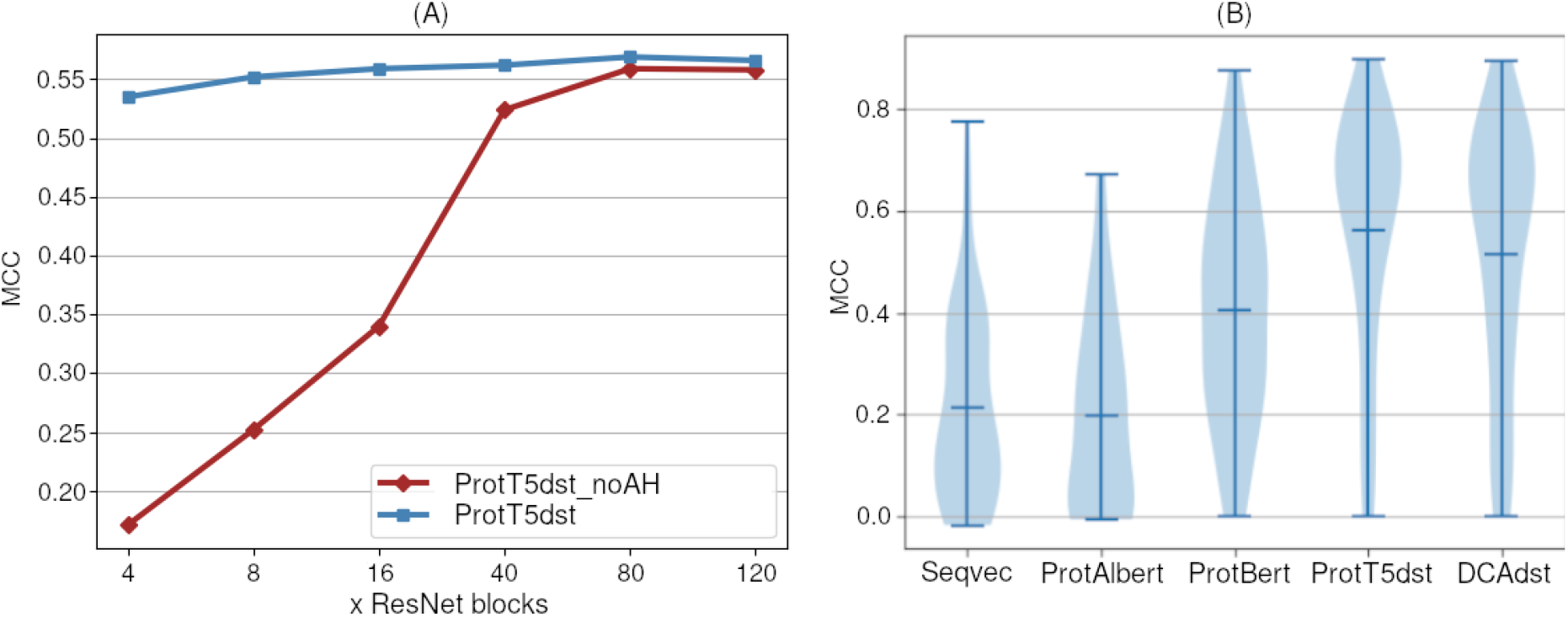
ProtT5 attention heads (AHs) best. Y-axes: Matthews Correlation Coefficient (MCC, Eqn. 4; for medium- and long-range contacts). <u>Panel A:</u> *SetValCASP12*; the x-axis gives the number of ResNet blocks; values on the left describe “shallow CNNs, i.e. those with fewer parameters (ranging from 235,306 for 4 blocks to 6,501,162 for 120 blocks). Both lines used the ProtT5 pLM: the upper blue line marks the method introduced here using the AHs, the lower red line embeddings without AHs. AHs already performed well with shallow architectures, while raw embeddings without AHs needed at least 40 residual blocks to reach MCC-levels above 0.5. <u>Panel B:</u> *SetValCASP12* + *SetTstCASP13*: the five violin plots for embeddings from four different pLMs (Elnaggar et al., 2021; Heinzinger et al., 2019) along with our in-house method using evolutionary couplings (DCAdst). The markers indicate highest, lowest and average MCC respectively, while the width – light-blue background cloud - shows the overall distribution.

### AHs clearly improved contact predictions

Even relatively shallow CNNs (relatively few free parameters) performed well when enriching the embeddings by using ProtT5 attention heads (AHs) rather than using embeddings without AHs (Fig. 2A). Smaller CNNs with 80 ResNet blocks (Fig. 1) even reached numerically higher MCCs than 50% larger CNNs with 120 ResNet blocks (Fig. 2A; difference not statistically significant). Nevertheless, all results given in the following were obtained for the less accurate version with 120 ResNet because we tested smaller CNNs after those results had been collected and decided to reduce energy-consumption not expecting significant improvements.

Comparing embeddings from different pLMs, Seqvec (based on ELMo (Peters et al., 2018)) and ProtAlbert (based on Albert (Lan et al., 2020), a leaner version of BERT (Devlin et al., 2019)) performed significantly worse than other transformers (Fig. 2B). Top were CNNs using the attention heads of ProtT5 (based on T5 (Raffel et al., 2020)) as input (Fig. 2B). Although this method never used any MSA for predicting inter-residue distances, it numerically even outperformed our in-house CNN dependent on evolutionary couplings (DCA, Fig. 2B).

Given that our method (ProtT5dst) used no MSA, and that it reached a similar average performance as our in-house CNN using evolutionary couplings (DCAdst), we expected embeddings to perform better for proteins from families with little sequence diversity (weak evolutionary coupling) and worse for those with large diversity (strong evolutionary coupling). Although, we observed evidence supporting this expectation (embeddings performed much better than evolutionary couplings for very small families), the attention head embeddings also performed better for some very large. We cannot explain this finding. One speculation is that very large families contain so much divergence in terms of sequence and structure that our embedding-based protein-specific predictions outperform the family-averaged predictions using evolutionary couplings. If so, at least the members of such large families most diverged in structure from the family-average might be predicted better without alignments (MSAs). As methods using evolutionary couplings benefit from immense diversity (Marks et al., 2012), simply constraining “too large families” might not remedy such a shortcoming of MSA-based solutions. If this speculation were partially correct, we have no data whether this would only affect the performance of some proteins (outliers) or of most (although most proteins might define the average, almost all might deviate substantially enough from the average).

### ProtT5dst with AHs not using MSAs reached Raptor-X relying on MSAs

For the CASP13 and CASP14 test sets (*SetTstCASP13+14*), we collected C-alpha contact predictions from Raptor-X, which is publicly available and performed well at CASP12 and CASP13 (Wang et al., 2017). We submitted only the original sequences instead of MSAs to allow the server to optimize its MSA. Given the database growth, Raptor-X most likely performed better when we tested it (May 2021) than at the CASP12/13 deadlines (summers 2016 and 2018, respectively). Although numerically, the supervised method ProtT5dst using AHs outperformed the version not using AHs (Table SOM2: ProtT5dst vs. ProtT5dst_noAH), this difference was not statistically significant within the 95% confidence interval, i.e. the values for the two models were within ±1.96 standard errors of each other. For medium-range contacts (between residue i,j with 12≤|i-j|≤23), AHs without MSAs numerically outperformed Raptor-X; for long-range contacts (|i-j|>23), largely the opposite was the case. However, none of those differences were statistically significant (Table SOM2: ProtT5dst vs. Raptor-X).

Comparing the embedding-based approach using AHs and no MSA (ProtT5dst) with the state-of-the-art Raptor-X using MSAs and post-processing for evolutionary couplings in detail, revealed that MSA-free predictions did perform better for very small families (Fig. 3A, darkest points usually above diagonal). For some proteins, (e.g. T0960-D2 and T0963-D2; Fig. 3A, top left), ProtT5dst correctly predicted distances while Raptor-X failed; for others, (e.g. T1049; Fig. 3A, bottom right), the opposite was the case.

**Fig. 3:**
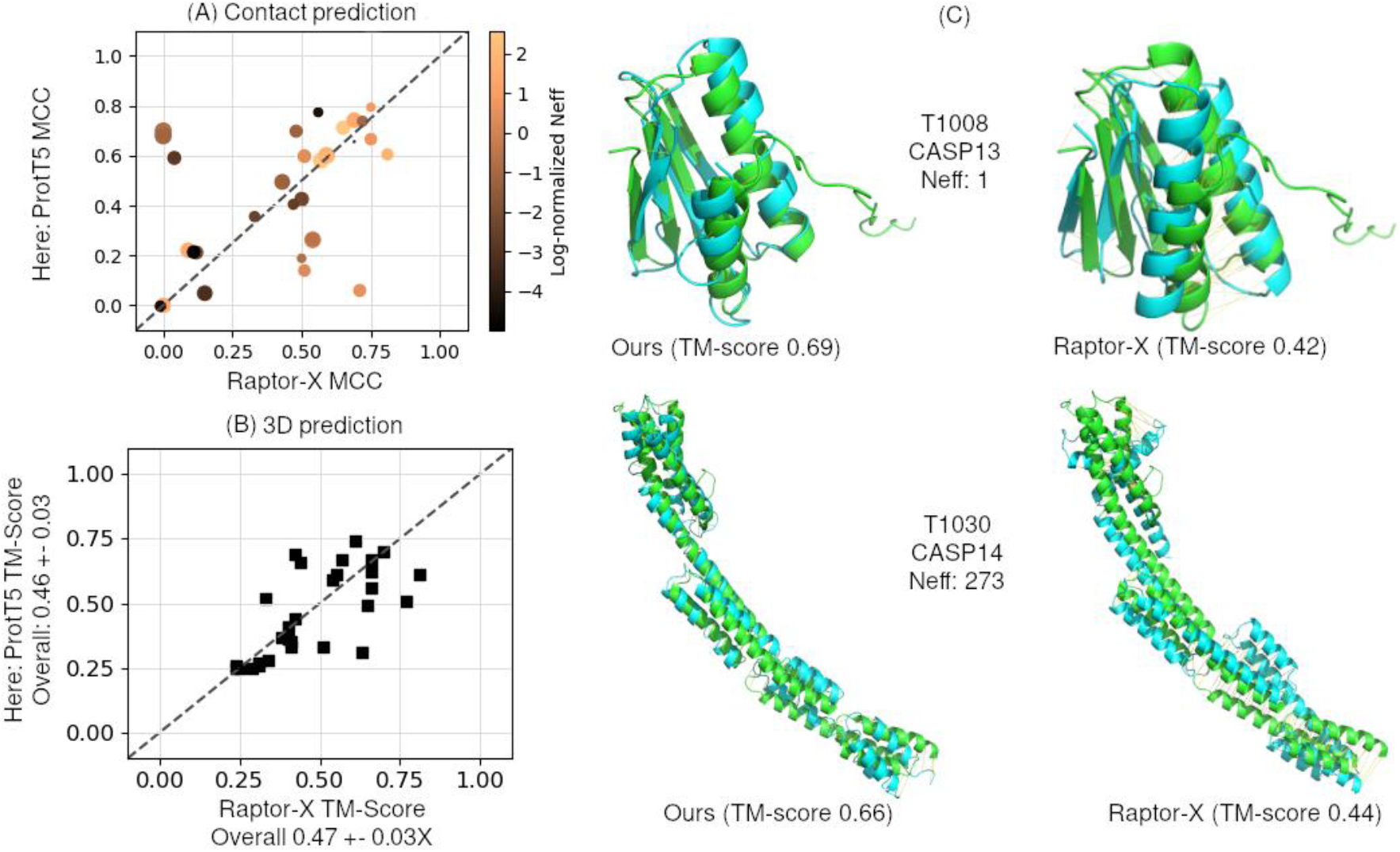
ProtT5dst beats Raptor-X for small families. *Data set:* on *SetTstCASP13* + *SetTstCASP14* (*methods):* ProtT5dst (as introduced here: using AHs without MSAs), and Raptor-X (Wang et al., 2017). *<u>Panel A</u>*: Per-protein comparison of MCCs (Eqn. 5) for medium- and long-range contact predictions. *<u>Panel B</u>*: 3D structure prediction performance: TM-align (Zhang & Skolnick, 2005) computed TM-scores for all predictions. Overall, the performance was similar for both methods. *<u>Panel C</u>*: Detailed comparison of 3D predictions vs. experiment for two proteins (T1008 and T1030; experiment: green, prediction: cyan). One protein gives an example for a small (T1008), the other for a large (T1030) family (T1008), and for both ProtT5 outperformed Raptor-X although overall the performance of these two was similar. For T1008 (CASP13), both predictions captured the overall fold correctly, but Raptor-X incorrectly swapped the two helices, reducing the TM-score from 0.69 (ProtT5dst) to 0.42. Similarly, for the longer protein T1030 (CASP14), Raptor-X misplaced several helices. Images from PyMol (Schrödinger & DeLano, 2021).

### Good 3D structure predictions

The trRosetta (Yang et al., 2020) pipeline with pyRosetta (Chaudhury et al., 2010) turned our predicted distance distributions (distograms) into 3D structure predictions. For the free-modeling and TBM-hard targets from CASP13 and CASP14, the similarity in terms of 2D predictions between ProtT5dst and Raptor-X remained essentially similar when using the predicted distances to predict 3D structure (Fig. 3A vs. 3B), with an average TM-score of 0.47 ± 0.03 for Raptor-X and of 0.46 ± 0.03 for ProtT5dst (Fig. 3B).

### Case study: beta-barrel gene duplication

All known transmembrane beta-barrel proteins, found in the outer membrane of Gram-negative bacteria, feature an even number of between 8 and 34 beta-strands (Georg, 2002). For instance OmpX from *Escherichia Coli* (Outer membrane protein protein X; Swiss-Prot identifier ompx_ecoli (Boutet et al., 2016)) has an 8-stranded beta-barrel. Gene *in vitro* duplication and selective removal of beta-hairpins produced new stable beta-barrel proteins, which folded *in vitro* with strand numbers between 8 and 16 (Arnold et al., 2007). Since none of the experimental structures of OmpX were included in any of our datasets, we could validate our model on its known structure (PDB identifier 1Q9F (Fernández et al., 2004)). ProtT5dst distance predictions refined through trRosetta (Yang et al., 2020) predicted the native OmpX structure accurately reaching a TM-score of 0.73 (Fig. 4 left). For three of the five engineered variants shown to fold in vitro (OmpX64c, OmpX66 and OmpX84), our predicted structures suggested a single larger barrel with 10 and 12 beta-strands (Fig. 4: three rightmost panels) that were confirmed experimentally (Arnold et al., 2007). As proof-of-principle, these results suggested that our approach can even succeed in reliably predicting structures of transmembrane proteins that are inherently difficult to predict by comparative modeling and other methods due to their under-representation in the PDB (Kloppmann et al., 2012; Pieper et al., 2013). The under-representation of membrane proteins in the PDB did not affect the pLMs underlying our predictions, because they only use sequence information and membrane proteins are likely not under-represented in UniProt (The UniProt Consortium, 2016).

**Fig. 4:**
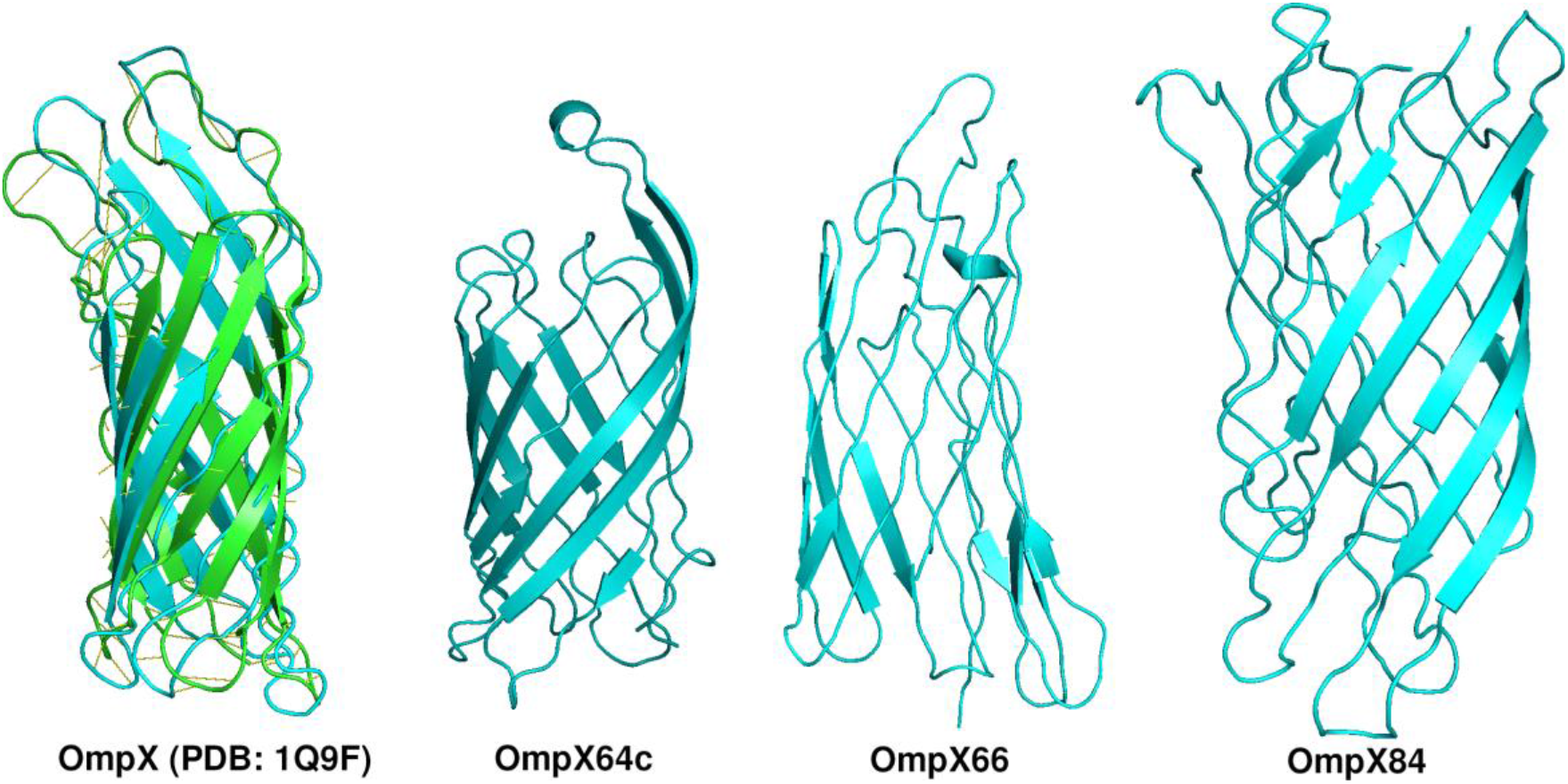
3D predictions for OmpX and 3 variants (OmpX64x, OmpX66 and OmpX84). The experimental structure is shown in green, predictions in cyan (images generated using PyMol). Prediction and experiment matched with a TM-score of 0.73 for the native protein of known structure. While the predictions for the protein-engineered sequence variants (OmpX64x, OmpX66, and OmpX84) suggested less compact structures, our predictions confirmed the experimental findings of larger single beta-barrels (Arnold et al., 2007).

### Saving computation time saves resources

Experimental high-resolution structures are so costly that good predictions are valuable even if they consume substantial amount of computing resources. The first compute-intensive tasks of state-of-the-art (SOTA) structure prediction methods is the generation of MSAs along with the processing of evolutionary couplings (Balakrishnan et al., 2011; Marks et al., 2011; Seemayer et al., 2014). Depending on hardware, alignment method, and sequence database, the average time needed to create MSAs varies substantially. For the 31 test proteins (SetTstCASP13 + SetTstCASP14), the total computation needed to infer distance distributions from the query sequence included using HHblits (Steinegger et al., 2019) on UniClust30 (2018_8) (Mirdita et al., 2016) to obtain MSAs, CCMpred (Seemayer et al., 2014) to generate couplings and running our in-house DCAdst prediction method. This took 212 minutes on an Intel Xeon Gold 6248 (100 GB RAM) and a single Nvidia Quadro RTX (46 GB VRAM) with all data on a local SSD. In contrast, using ProtT5dst on the same hardware, pre-dictions completed in only 2 minutes and 11 seconds, corresponding to an almost 100-fold speed-up. The runtime measures included loading our pre-trained models, amounting to a one-time cost of ∼25 seconds regardless of the number of proteins predicted. We computed predictions for almost the entire human proteome (proteins with <3000 residues due to GPU memory limits) in about eight days, using the same hardware. Obviously, the measures did not consider the time for pLM pre-training (ProtT5) because that method had been made available before we started and has not been tailored in any way to predict protein structure.

### Distance predictions for almost all human proteins

Just when we had completed of inter-residue distance predictions for all human proteins below 3k residues, the 3D predictions from *AlphaFold 2* were made available for all those proteins (Tunyasuvunakool et al., 2021). Does any reason remain to have our resource in parallel? Although we are currently unsure how we will proceed, i.e. whether or not we will invest any more computing resource to grow our database of 2D predictions, we clearly see an advantage in the simplicity of our resource. Although source code and Colab notebook are available for AlphaFold 2 (Tunyasuvunakool et al., 2021), applying those on your protein might be more challenging than applying the simple DCAdst 2D-distance predictions to the protein of your interest (in case that is not already contained in the growing database of AlphaFold 2 predictions). It remains to be investigated, to which extent more protein-specific vs. more family-averaged predictions will matter. Will predictions based on simple embeddings be more useful for protein design (Wu et al., 2021) than those based on MSAs, or will the best use both approaches? Too early to tell.

## Conclusions

We showed that 2D inter-distance predictions based on embeddings derived from single protein sequences improved significantly over recent years and now rival the performance of co-evolution methods. While our approach does not improve SOTA performance yet, the vast reduction in inference time without sacrificing prediction accuracy provides a crucial practical advantage. Even more importantly, our approach offers, for the first time, an accurate protein structure prediction based on single protein sequences that is competitive to family-centric approaches that rely on diverse MSAs. Since structure predictions can be obtained in mere seconds, our method could easily provide the basis for high-throughput analysis of protein structure predictions, such as *in silico* structure mutation. It is likely that approaches using embeddings and co-evolution information will co-exists in the future and might provide mutual benefits. In the future, we will be investigating the feasibility of MSA refinement using our method by filtering aligned sequences by predicted structural differences.

## Availability

Pre-trained models and the source code for the prediction pipeline are available at https://github.com/kWeissenow/ProtT5dst.

Our predictions for all human proteins (<3000 residues) are stored at https://rostlab.org/∼conpred/ProtT5dst/pred_all_human/ (work in progress).

## Acknowledgements

The authors thank primarily Tim Karl (TUM) for invaluable help with hard- and software and Inga Weise (TUM) for support with many other aspects of this work. We thank Nir Ben-Tal for helpful suggestions and inspiration for the OmpX case study. Thanks to Jinbo Xu and his co-developers of Raptor-X for making their method available; thanks to Jianyi Yang and his co-developers for publishing the trRosetta source code. This work was supported by a grant from the Alexander von Humboldt foundation (BMBF), and by a grant from the German Research Foundation (Deutsche ForschungsGemeinschaft; DFG–GZ: RO1320/4–1). We gratefully acknowledge the support of NVIDIA with the donation of a Titan GPU used for the development phase. Furthermore, the Rostlab gladly acknowledges support from Google Cloud and Google Cloud Research Credits program to fund the earlier stages of this project under the Covid19 HPC Consortium grant. Last not least, thanks to all who make their experimental data publicly available and all those who maintain such databases, in particular to Steve Burley and his team at the PDB.

## Abbreviations used

2D: two-dimensional
2D structure: inter-residue distances/contacts
3D: three-dimensional
3D structure: coordinates of atoms in a protein structure
APC: average product correction
CASP: Critical Assessment of protein Structure Prediction
CNN: convolutional neural network
DCA: direct coupling analysis
DL: Deep Learning
LM: Language Model
LR: logistic regression
MSA: multiple sequence alignment
pLM: protein Language Model
SOTA: state-of-the-art

