## Supporting Online Material for "Protein language model embeddings for fast, accurate, alignment-free protein structure prediction"

### Supporting Online Material (SOM) for:
### Protein language model embeddings for fast, accurate, alignment-free protein structure prediction

**Konstantin Weißenow[1,2,*], Michael Heinzinger[1,2] & Burkhard Rost [1,3]**

1 TUM (Technical University of Munich) Department of Informatics, Bioinformatics & Computational Biology - i12, Boltzmannstr. 3, 85748 Garching/Munich, Germany
2 TUM Graduate School, Center of Doctoral Studies in Informatics and its Applications (CeDoSIA), Boltzmannstr. 11, 85748 Garching, Germany
3 Institute for Advanced Study (TUM-IAS), Lichtenbergstr. 2a, 85748 Garching/Munich, Germany & TUM School of Life Sciences Weihenstephan (WZW), Alte Akademie 8, Freising, Germany & Columbia University

#### Table of Contents for SOM

#### Short description of SOM

For brevity, we omitted a full visualization of our ResNet architecture in the main manuscript, which is shown here in Fig. SOM_1 in more detail.

Table SOM_1 shows our results with logistic regression on attention heads of our ProtT5 pLM, highlighting that a reduction from all 768 heads down to the 100 most informative still yields comparable performance.

Finally, in table SOM_2 we contrast our results using ProtT5 embeddings and attention heads (as well as an older pLM model for comparison) with Raptor-X (Wang et al., 2017).

### Material

**Fig. SOM_1: Architecture of ProtT5dst deep dilated residual CNN.**

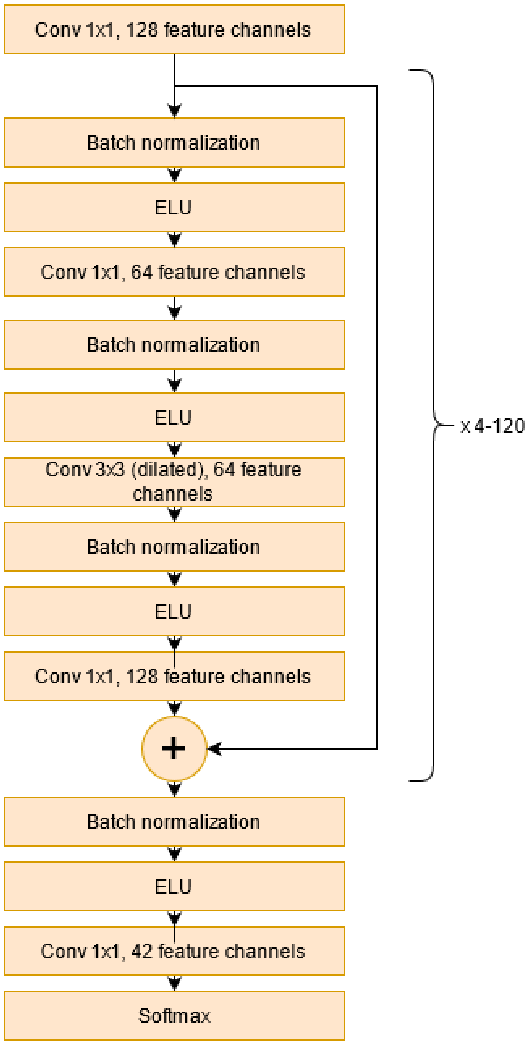

**Fig. SOM_1: Architecture of ProtT5dst deep dilated residual CNN.** We use ResNet blocks stacked 4 to 120 times depending on input type (see Methods). Similar to AlphaFold (Senior et al., 2020), each residual block contains three convolution operations, the first and last being down- and upscaling layers, while the central 3x3 convolution cycles through dilation factors 1 to 8 in each successive block.

**Table SOM_1: Logistic regression results on ProtT5 attention heads.**

| | *Precision %* | *Recall %* | *F-Score %* | *MCC* | *MCC (long-range)* |
|---|---|---|---|---|---|
| *All 768 heads* | 58 ± 6 | 15 ± 3 | 22 ± 3 | 0.27 ± 0.04 | 0.23 ± 0.04 |
| *All 768 heads + sym* | 60 ± 6 | 17 ± 3 | 25 ± 3 | 0.30 ± 0.04 | 0.25 ± 0.04 |
| *All 768 heads + sym + APC* | 57 ± 6 | 18 ± 3 | 26 ± 3 | 0.30 ± 0.04 | 0.25 ± 0.04 |
| *Top-50 + sym + APC* | 52 ± 6 | 15 ± 3 | 23 ± 3 | 0.26 ± 0.04 | 0.24 ± 0.04 |
| *Top-100 + sym +APC* | 56 ± 6 | 17 ± 3 | 25 ± 3 | 0.29 ± 0.04 | 0.24 ± 0.04 |
| *Top-120 + sym + APC* | 56 ± 6 | 17 ± 3 | 25 ± 3 | 0.29 ± 0.04 | 0.25 ± 0.04 |

* Logistic regression results based on attention heads from the ProtT5 pLM for 200 randomly selected training samples assessed on S*etValCASP12* validation set. Methods (rows): The first three rows show results based on all 768 attention heads directly, with symmetry (+sym) and with symmetry and average product correction (+APC). The lower 3 rows show the performance of the Top-50, Top-100 and Top-120 most informative attention heads. Performance measures (columns): The ± values indicate ±1.96 standard errors, i.e. 95% confidence interval (CI95; Eqn. 7). The 100 most informative attention heads suffice to reach baseline performance (within the standard error).

**Table SOM_2: Performance of embeddings similar to RaptorX ***

| | *Medium-range* | | | *Long-range* | | | Top N Precision with N= | | |
|---|---|---|---|---|---|---|---|---|---|
| | *P (Eqn. 2)* | *R (Eqn. 3)* | *MCC (Eqn. 5)* | *P (Eqn. 2)* | *R (Eqn. 3)* | *MCC (Eqn. 5)* | L/1 | L/2 | L/10 |
| *ProtBert* | 55 ± 4 | 20 ± 3 | 0.30 ± 0.03 | 25 ± 6 | 8 ± 3 | 0.12 ± 0.03 | 24 ± 5 | 27 ± 6 | 30 ± 6 |
| *ProtT5dst_noAH* | 63 ± 3 | **34** ± **4** | 0.43 ± 0.04 | 28 ± 5 | 16 ± 4 | 0.20 ± 0.04 | 31 ±5 | 38 ± 6 | 42 ± 7 |
| *ProtT5dst* | **67** ± **4** | 33 ± 4 | **0.43** ± **0.04** | 36 ± 7 | **18** ± **4** | 0.24 ± 0.05 | 35 ± 5 | 42 ± 6 | 51 ± 7 |
| *Raptor-X* | 66 ± 5 | 31 ± 4 | 0.41 ± 0.04 | **45** ± **7** | **18 ± 3** | **0.26 ± 0.04** | **40** ± **5** | **48** ± **7** | **59** ± **7** |

* Data sets: SetTstCASP13 and SetTstCASP14 free modeling and TBM-hard targets; Medium-range: sequence distance between 4 and 23 residues; Long-range: sequence distance >23 residues; Top N: Top-L precision (see methods) evaluated for L/1, L/2 and L/10; Measures: Precision (P), Recall (R), and Matthew correlation coefficient (MCC); ± values indicate ±1.96 standard errors, i.e. 95% confidence interval (CI95, Eqn. 7). Methods: we compare performance of our predictors using 1D embeddings (ProtT5dst_noAH) and attention heads

(ProtT5dst) as well as our previous best-performing language model (ProtBert) with Raptor-X (Wang *et al.*, 2017). ; bold face: numerically highest method in each column in bold.
